# The Cell Adhesion Molecule TMIGD1 Binds to Moesin and Regulates Tubulin Acetylation and Cell Migration

**DOI:** 10.1101/2021.08.08.455580

**Authors:** Nader Rahimi, Rachel X-Y Ho, Kevin Brown Chandler, Kyle Oliver Corcino De La Cena, Razie Amraei, Ashley J Mitchel, Nels Engblom, Catherine E Costello

**Author notes:** **Corresponding Authors:** Nader Rahimi, and Catherine E Costello,.

## Abstract

**Background:** The cell adhesion molecule transmembrane and immunoglobulin (Ig) domain containing1 (TMIGD1) is a novel tumor suppressor that plays important roles in regulating cell-cell adhesion, cell proliferation and cell cycle. However, the mechanisms of TMIGD1 signaling are not yet fully elucidated.

**Results:** TMIGD1 binds to the ERM family proteins moesin and ezrin, and an evolutionarily conserved RRKK motif on the carboxyl terminus of TMIGD1 mediates the interaction of TMIGD1 with the N-terminal ERM domains of moesin and ezrin. TMIGD1 governs the apical localization of moesin and ezrin, as the loss of TMIGD1 in mice altered apical localization of moesin and ezrin in epithelial cells. In cell culture, TMIGD1 inhibited moesin-induced filopodia-like protrusions and cell migration. More importantly, TMIGD1 stimulated the Lysine (K40) acetylation of α-tubulin and promoted mitotic spindle organization and CRISPR/Cas9-mediated knockout of moesin impaired the TMIGD1-mediated acetylation of α-tubulin and filamentous (F)-actin organization.

**Conclusions:** TMIGD1 binds to moesin and ezrin, and regulates their cellular localization. Moesin plays critical roles in TMIGD1-dependent acetylation of α-tubulin, mitotic spindle organization and cell migration. Our findings offer a molecular framework for understanding the complex functional interplay between TMIGD1 and the ERM family proteins in the regulation of cell adhesion and mitotic spindle assembly, and have wide-ranging implications in physiological and pathological processes such as cancer progression.

## Introduction

Altered cell-cell adhesion and disruption of apico-basal polarity are common features of all carcinomas [14,40]. Tumor suppressor genes such as *VHL* and *APC* that are frequently inactivated in human renal and colon cancers, are also key regulators of cell adhesion and polarity[13,23]. First discovered as a regulator of cell-cell adhesion and cell morphology, the cell adhesion molecule transmembrane and immunoglobulin (Ig) domain containing-1 (TMIGD1) is predominantly expressed in kidney and intestinal epithelial cells and protects renal epithelial cells from oxidative cell injury, thus promoting cell survival [2]. Emerging evidence on the role of TMIGD1 in human cancers points to TMIGD1 as a novel tumor suppressor. TMIGD1 is downregulated in human renal and colon cancers [10,27]. Re-expression of TMIGD1 in renal and colon cancer cell lines inhibits cell proliferation and induces G2/M cell cycle checkpoint arrest [10,27]. Recent studies have revealed that loss of TMIGD1 in mice significantly impairs intestinal epithelium brush border membrane junctional polarity and maturation, resulting in the development of adenomas in small intestine and colon [10]. TMIGD1 inhibits tumor cell proliferation and cell cycle arrest at the G2/M phase through regulating expression of p21^CIP1^ (cyclin-dependent kinase inhibitor 1), and p27^KIP1^ (cyclin-dependent kinase inhibitor 1B)[10]. However, the specific mechanisms by which TMIGD1 elicits these effects, particularly at the level of protein-protein interaction remain unknown. TMIGD1 is a transmembrane glycoprotein that consists of an extracellular domain with two Ig domains, a single transmembrane domain, and a highly conserved short intracellular domain enriched in the positively charged amino acids Lysine (K) and Arginine (R)[2], with a potential to recruit signaling proteins to TMIGD1 [2]. Ezrin, radixin and moesin (ERM), three highly similar proteins are members of the FERM (4.1-band ERM) superfamily, which are central for linking the actin cytoskeleton to the cell membrane and are major regulators of specialized membrane domains, including apical microvilli[22,39], lamellipodia and filopodia[4,21]. Due to their function as cytoskeletal linkers, ERM family proteins play essential roles in diverse cellular processes ranging from cell-cell adhesion and cell migration to cell proliferation[8], and thus play significant roles in the metastatic progression of human cancers. Furthermore, the FERM protein moesin is known to bind to microtubules (MTs) and mediate the association of actin filaments, which is required for regulating spindle organization during mitosis [36,42]. MTs are cytoskeletal filaments composed of heterodimers of α- and β-tubulin subunits and are critically important for chromosomal segregation, intracellular transport, cell division, cell motility and cell morphogenesis[30]. Post translational acetylation of tubulin stabilizes MTs and plays a central role in its dynamic features and cellular functions, a process exploited by anti-tumor agents such as Taxol to promote mitotic arrest and cell death[43]. Here, we present evidence that TMIGD1 binds to ERM family proteins (moesin and ezrin), regulates the stability of microtubules and modulates cell migration in renal cancer cells, implicating TMIGD1 as a potential cancer therapeutic target.

## Materials and Methods

### Plasmids and Antibodies

Moesin-GFP, ezrin-GFP, GST-FERM-moesin, GST-C-ERMAD-moesin, and LifeAct7 plasmids were purchased from Addgene. Moesin sgRNA was purchased from Dharmacon. Construction and cloning of TMIGD1 into retroviral, pMSCV.puro vector was previously described [2,27]. GST-TMIGD1 and GST-4A-TMIGD1 were generated by standard PCR amplification of the cytoplasmic region of TMIGD1 and cloned into pGEX-4T-2 vector. Plasmids were expressed in *E*.*coli* and GST-fusion proteins were purified from E.coli via glutathione Sepharose Fast-flow resin per manufacturer’s recommendations. Polyclonal rabbit anti-α-tubulin antibody, polyclonal rabbit-anti-acetyl (K40)-tubulin antibody and anti-Myc antibody, and anti-GFP antibody all were purchased from Cell Signaling Technology Inc., (Danvers, MA). Development and characterization of the polyclonal anti-TMIGD1 antibody used in these studies has been described previously [2].

### Cell culture

HEK-293, RKO, 786-0 cells were each maintained in Dulbecco’s modified Eagle medium (DMEM) supplemented with 10% fetal bovine serum (FBS), L-glutamine (2 mM), penicillin (50 units/ml) and streptomycin (50 mg/ml).

### Animal Studies

CRISPR/Cas9-mediated TMIGD1 knockout mice were developed in our laboratory [10]. PFA fixed tissues were stained for moesin, ezrin or for anther protein of interest, as described in the figure legends.

### Retrovirus production and transfection

pMSCV.puro vector containing TMIGD1 or another cDNA of interest was transfected into 293-GPG cells, and viral supernatants were collected for five days as previously described [32] and viral supernatants were used to generate stable RKO or HEK-293 cell lines expressing TMIGD1 or empty vector (EV) as a control. TMIGD1 expressing cells were selected with puromycin. In some experiments as described in the figure legends, these cell lines were transiently transfected with moesin-GFP, ezrin-GFP or other plasmids via PEI (polyethylenimine). After 48 hours, cells were lysed and subjected to co-immunoprecipitation or western blotting as described in the figure legends.

### Immunoprecipitation and western blot analysis

Cells were grown in 10-cm culture dishes until 80 to 90% confluence. Cells were lysed, and normalized whole-cell lysates were subjected to immunoprecipitation by incubation with appropriate antibodies as shown in the figure legends. Immuno-complexes were captured by incubation with either protein A-Sepharose or protein G-agarose beads. After release via boiling the samples for 5 min at 95 °C, the immunoprecipitated proteins were subjected to western blot analysis. Occasionally, membranes were stripped by incubating them in a stripping buffer containing 6.25 mM Tris-HCl, pH 6.8, 2% SDS, and 100 mM β-mercaptoethanol at 50 °C for 30 min, washed in Western Rinse buffer (20 mM Tris and 150 mM NaCl), and re-probed with the antibody of interest. The blots were scanned and subsequently quantified using Image J (NIH).

### GST-pulldown Assay

*In vitro* GST fusion protein binding experiments were performed as described previously [26]. Briefly, equal numbers of cells expressing TMIGD1 or other proteins, as described in the figure legends were grown to 90% confluence. Cells were lysed in ice-cold lysis buffer supplemented with 2 mM Na3VO4 and a protease inhibitor cocktail. Equal amounts of the appropriate immobilized GST fusion proteins were incubated with normalized whole-cell lysates by rocking for 3 h at 4 °C. The beads were washed in the presence of protease inhibitors, and proteins were eluted and analyzed by western blotting using the appropriate antibody, as described in the figure legends.

### Liquid chromatography–tandem mass spectrometry (LC-MS/MS)

Agarose resins with GST-TMIGD1 bound via glutathione were incubated with whole cell lysate derived from 786-0 cells. Captured proteins were resolved on SDS-PAGE. Bands of interest were excised from the gel, and individual bands were subjected to trypsin digestion (at 37 °C overnight in 50 mM ammonium bicarbonate). Peptides were separated and analyzed via a 6550 Q-TOF MS equipped with a 1200 series nanoflow HPLC-Chip ESI source fitted with an HPLC-Chip consisting of a 360-nl trapping column and a 150 mm × 75 μm analytical column, both packed with Polaris C18-A 3-μm material (all from Agilent Corp.). After injection of the sample onto the trapping column, the column was washed at a rate of 2 μl/min with 2% acetonitrile and 0.1% formic acid in water for 4 min. Peptides were then separated on the analytical column at a flow rate of 0.3 μl/min using a gradient from 2% to 40% acetonitrile with 0.1% formic acid over 25 min. The 6550 Q-TOF mass spectrometer was operated in the positive-ion mode using the high-resolution, extended dynamic range (2 GHz) setting. The instrument was operated in data-dependent mode; the 20 most abundant ions were selected for MS/MS. MS spectra were recorded over the range *m/z* 285–1700, and MS2 spectra were recorded from *m/z* 50–3000. The ion source gas temperature was set to 225 °C, and the flow was set at 13 l/min, with a capillary voltage of 2100 V. Precursors ≥5000 counts and charge states ≥2 were selected for fragmentation, and the collision energy was set according to the equation y = mx + b, with y being the collision energy, slope m = 3.6, x representing the charge state, and the offset b = −4.8 for charge states 3 and 4. For charge state 2 peptides, slope m = 3.1, and the offset b = 1. Spectra were recorded in the centroid mode. MS/MS spectra were searched using a local copy of Mascot (www.matrixscience.com) using the Uniprot database of human proteins. The selected parameters required a minimum of two peptide matches per protein, with minimum probabilities of 95% at the peptide level.

### Immunofluorescence Microscopy

Cells expressing TMIGD1 alone or together with moesin were seeded onto coverslips and grown overnight in 60-mm plates to 80-100% confluence. The cells were washed once with PBS and fixed with freshly prepared 4% paraformaldehyde for 15 min at room temperature. After being washed three more times with PBS, the cells were permeabilized with 0.25% Triton X-100 in Western rinse for 10 min at room temperature and then washed three times with PBS. For staining with anti-*N*-acetyl tubulin or total tubulin, the cells were blocked with (1:1) BSA in Western Rinse buffer for 1 h and washed once in PBS, followed with incubation with the selected antibody, as indicated in the figure legends, for 1 h and then detected with a FITC-conjugated secondary antibody. The coverslips were mounted in Vectashield mounting medium with DAPI onto glass microscope slides. The slides were analyzed under a fluorescence microscope.

### F-actin stress orientation quantification

786-0 cells expressing TMIGD1 alone or with other constructs were stained for actin, as described above, and visualized using an inverted epifluorescence microscope. From each image, five different regions were chosen randomly for evaluation. F-actin orientation (anisotropy) and expression were quantified using the open source plugin Fibriltool for Image J as described [15].

### Cell Migration assay and filopodia assessment

Cell migration was carried out using Boyden Chamber assay as previously described [2,33]. To assess filopodia formation, RKO cells expressing control vector (EV) or TMIGD1 were transfected with moesin-GFP or control vector, GFP. Cells were fixed and pictures were taken under Nikon Deconvolution microscope equipped with camera. Quantification of filopodia was carried out using an open source software, filopodyan [41].

### Statistical analyses

Experimental data were subjected to Student t-test or One-way analysis of variance, where appropriate, with representation of at least three independent experiments. p<0.05 was considered significant, except where indicated otherwise in the figure legends.

## Results

### Identification of Moesin as a TMIGD1 Binding Protein

To identify TMIGD1 interacting proteins in renal cancer cells, we generated a recombinant GST-fusion protein encompassing the cytoplasmic domain of human TMIGD1 (**Figure 1A**). We incubated whole cell lysate derived from 786-0 renal carcinoma cells with the agarose resin bound with GST-TMIGD1 and analyzed the GST-TMIGD1 captured proteins via liquid chromatography–tandem mass spectrometry (LC-MS/MS). Moesin was one of the distinct proteins identified in these GST-TMIGD1 pull-downs (**Figure 1B**). We further validated the binding of TMIGD1 with moesin in HEK-293 cells ectopically expressing moesin-GFP via this same GST-TMIGD1 pull-down assay; moesin-GFP was selectively pulled-down with GST-TMIGD1 (**Figure 1C**). In an alternative approach, we used 786-0 cells expressing either empty vector (EV) or TMIGD1 and determined the binding of endogenously expressed moesin with TMIGD1 in 786-0 cells via immunoprecipitation with the anti-TMIGD1 antibody. This assay also showed that moesin binds to TMIGD1 (**Figure 1D**). Expression of endogenous TMIGD1 in 786-0 cells is very low/undetectable as previously reported[27].

**Figure 1:**
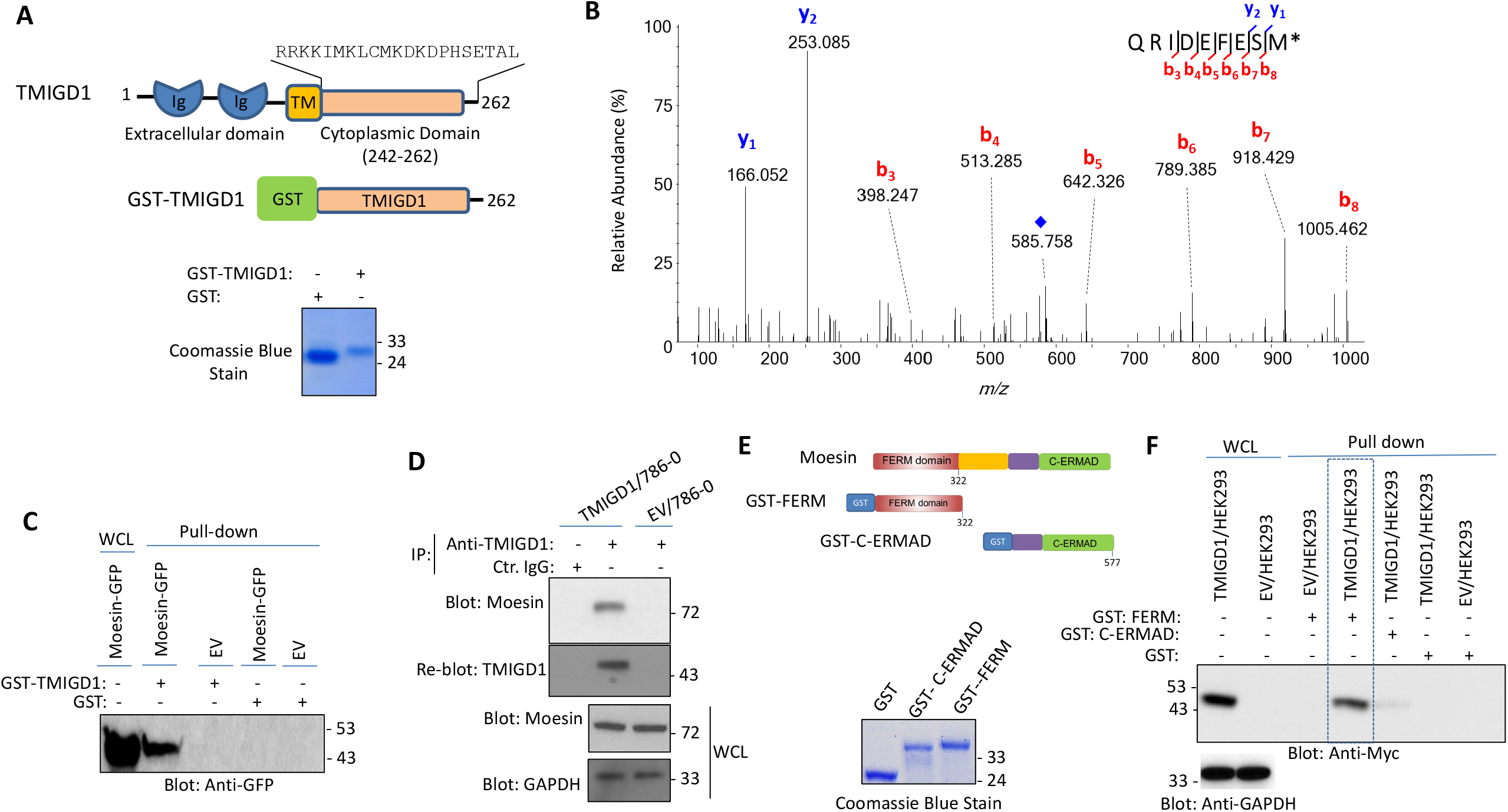
Identification of moesin as a TMIGD1 binding protein: (**A**) Schematic of GST-TMIGD1 cytoplasmic domain and its expression. (**B**) Collision-induced dissociation (CID) MS/MS spectrum assigned to the C-terminal Moesin peptide QRIDEFESM[O] (precursor [M + 2H]^2+^ *m/z* 585.578, one of four high-confidence Moesin peptide matches for analysis of the tryptic digestion products of xx. *Indicates oxidation of methionine; blue diamond indicates the residual precursor ion. (**C**) Whole cell lysates from HEK-293 cells expressing GFP tagged moesin or control vector (EV) were incubated with GST or GST-TMIGD1 coated beads. The captured proteins were released and resolved on SDS-PAGE followed by Western blotting using an anti-GFP antibody. (**D**) Whole cell lysates from 786-0 cells expressing EV or TMIGD1 were incubated with anti-TMIGD1 antibody and immunoprecipitated proteins were detected via Western blot analysis using an anti-Moesin antibody. The same membrane was blotted for TMIGD1. Whole cell lysates (WCL) from the same group were blotted for total Moesin or GAPDH. (**E**) Schematic of GST Moesin constructs encompassing the N-terminal GST-FERM or the C-terminal GST-C-ERMAD domains. Coomassie Blue stain of GST, GST-FERM and GST-C-ERMAD in Whole Cell Lysate (WCL) or pulled-down proteins. (**F**) Whole cell lysates from HEK-293 cells expressing EV or TMIGD1-Myc were incubated with GST, GST-FERM or GST-C-ERMAD followed by Western blot analysis using anti-Myc antibody.

Moesin is composed of two major domains: the N-terminal FERM domain and the C-terminal ERMAD domain (**Figure 1E**). As such, we sought to determine which of the two domains of moesin was involved in interacting with TMIGD1. To answer this question, we generated GST-FERM and GST-ERMAD fusion proteins (**Figure 1E**) and evaluated their ability to interact with TMIGD1 in a GST-pulldown assay. The result showed that GST-FERM moesin selectively binds to TMIGD1 (**Figure 1F**). Also noted, was a faint protein band detected with GST-ERMAD-moesin, potentially suggesting a weak interaction with TMIGD1 (**Figure 1F**). The data demonstrate that moesin primarily interacts with TMIGD1 via its N-terminal FERM domain.

ERM proteins are highly conserved, possessing more than 75% amino acid homology within the common FERM domain[12]. The N-terminal FERM domain is known to recognize positively charged amino acid clusters (*i*.*e*., lysine/arginine)[44]. The amino acid sequence of moesin and ezrin are shown (**S. Figure 1**). Notably, the cytoplasmic domain of TMIGD1 contains multiple positively charged amino acid clusters, which are conserved between human and mouse TMIGD1 (**Figure 2A**). These positively charged amino acid clusters present on TMIGD1 have similarity to those found on CD44, CD43 and ICAM2, which are known to mediate the binding of ERM proteins[44] (**Figure 2A**). These residues on TMIGD1 are thus likely to be involved in the recognition of moesin. To investigate this possibility, we generated GST-fusion TMIGD1 and GST-4A-TMIGD1 proteins in which two arginine (R, 242 &243) and two lysine (K, 244 &245) residues were mutated to alanine (A) (**Figure 2B**). The purified recombinant proteins were used to evaluate their interaction with moesin-GFP and ezrin-GFP expressed in HEK-293 cells (**Figure 2C**). As anticipated, removal of the four positively charged residues (242-245) abolished the interaction of moesin and ezrin with TMIGD1 (**Figure 2D**). Our data demonstrate that TMIGD1 interaction with ERM family proteins is established via the RRKK (242-245) motif present in the conserved C-terminus of TMIGD1.

**Figure 2:**
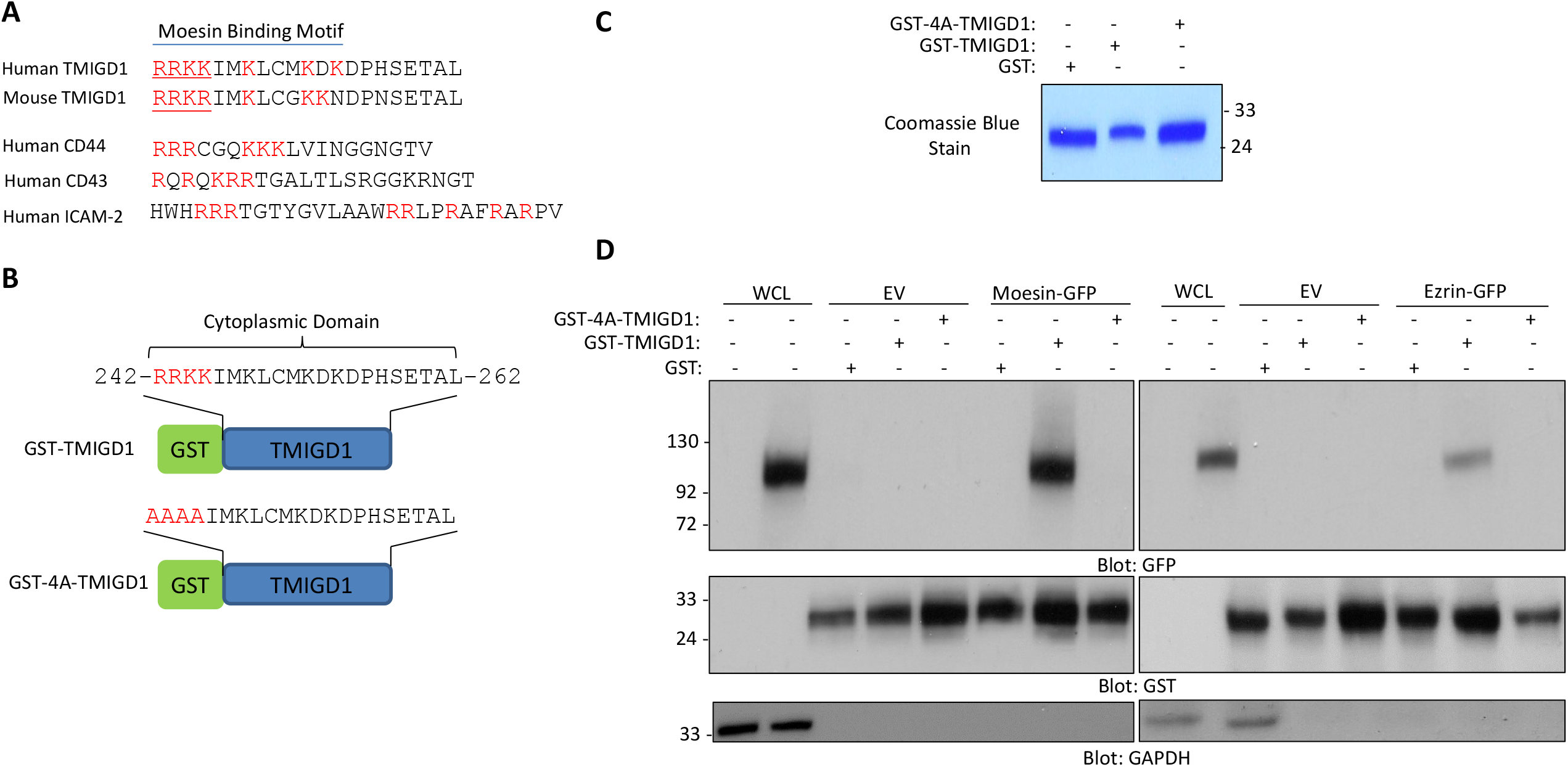
Conserved RRKK motif mediates TMIGD1 binding with ERM proteins Moesin and Ezrin. **(A**) Amino acid sequence homology of human and mouse TMIGD1 and sequence of the known Moesin binding receptors. (**B**) Schematic of wild type GST-TMIGD1 and GST-RRKK motif mutant TMIGD1 (4A-TMIGD1). (**C**) Equal amount of purified recombinant Glutathione S-transferase (GST), GST-TMIGD1 and 4A-TMIGD1 fusion proteins were resolved on SDS-PAGE followed by Coommassie blue staining. (**D**) Whole cell lysates derived from HEK-293 cells expressing empty vector (EV), GFP-tagged moesin or GFP-tagged ezrin were incubated with GST, GST-TMIGD1 or GST-4A-TMIGD1 followed by western blot analysis using an anti-GFP antibody. The same membranes were blotted for GST or GAPDH.

### Loss of TMIGD1 in Mice Alters Apical Localization of ERM proteins Moesin and Ezrin in Epithelial cells

ERM proteins are commonly localized at the apical membranes of polarized epithelial cells where they link filamentous (F)-actin to plasma membrane proteins [28,34]. Similar to the ERM proteins, TMIGD1 is also localized at the apical region of renal and intestinal epithelial cells [2,10] (**Figure 3A**). Our recent study demonstrated that loss of TMIGD1 in mice alters intestinal apical membrane organization [10]. Therefore, we asked whether the loss of TMIGD1 affects localization of the ERM proteins to apical membranes. Immunofluorescence staining of kidney tissues from wild type mice (TMIGD1+/+) and TMIGD1 KO mice (TMIGD1-/-) revealed that, while moesin is largely localized to the apical membranes of renal epithelial cells in the TMIGD1+/+ mouse, this apical localization and organization was decreased in TMIGD1-/- mice (**Figure 3B**). Genotypic analysis of TMIGD1 KO mouse used in this study is shown (**S. Figure 2**). Additionally, we stained mouse intestinal tissues for ezrin, which is highly expressed in intestinal epithelial cells and required for apical membrane organization [34]. Just as we had observed for moesin in renal epithelial cells, ezrin localization at the apical membranes in TMIGD1-/- mice was also significantly impaired (**Figure 3C**).

**Figure 3.**
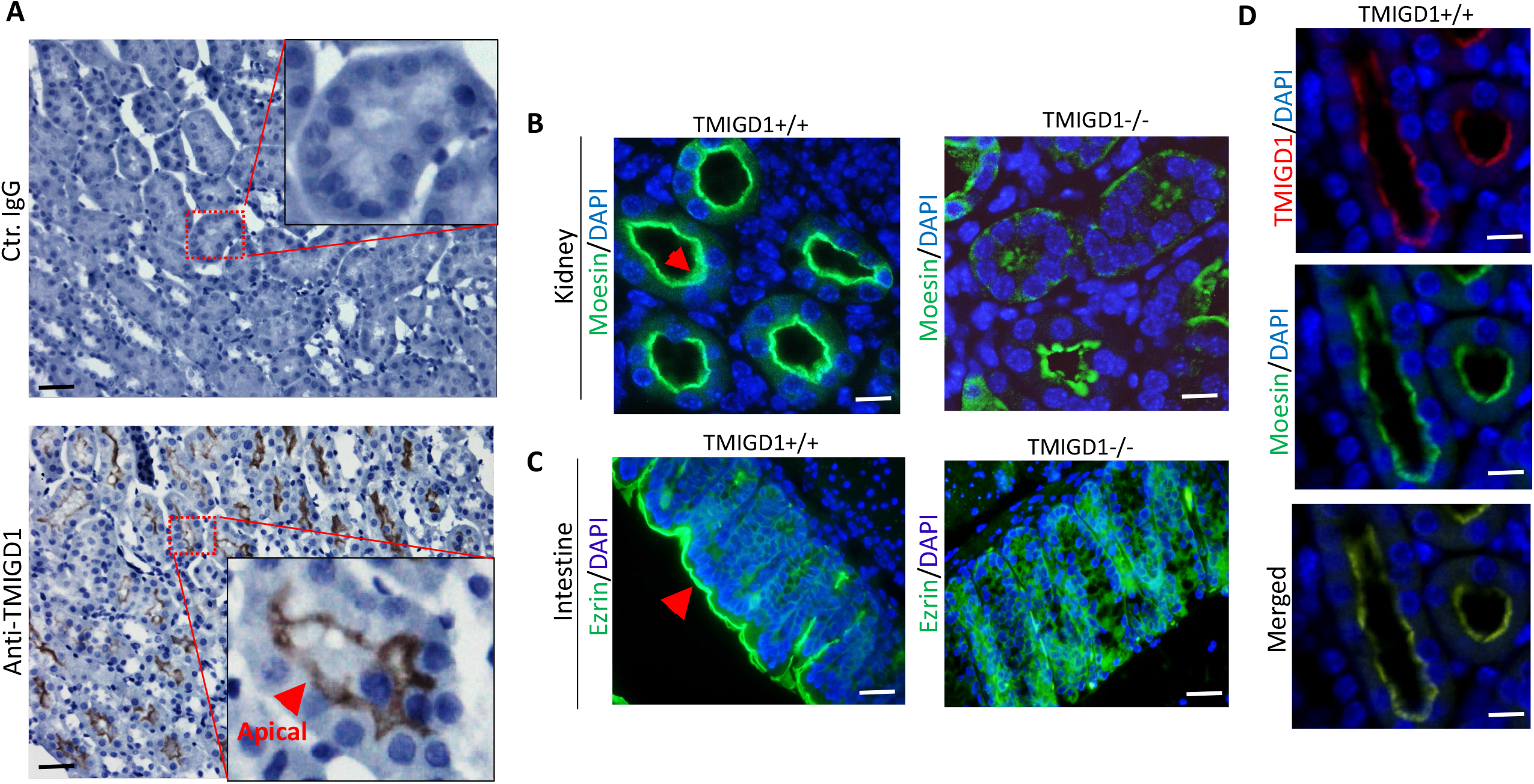
Loss of TMIGD1 in mice interferes with Moesin and Ezrin apical membrane localization. (**A**) Immunohistochemistry staining of human kidney tissues with control antibody or with anti-TMIGD1 antibody. Image magnification 50 µM. Arrow points to apical membrane localization of TMIGD1. (**B**) Immunofluorescence staining of wild type kidney and intestine tissues from (TMIGD1+/+) or TMIGD1 knockout mice (TMIGD1-/-) stained with antibodies against moesin and ezrin, respectively. Red arrows point to the apical membrane. Image magnification 50 µM. (**C**) Immunofluorescence staining of wild type mice intestine tissue co-stained with moesin and TMIGD1. Image magnification 50 µM.

Considering the observed binding of TMIGD1 with moesin, we also examined the co-localization of TMIGD1 with moesin. Immunofluorescence staining of mouse kidney tissue showed that TMIGD-1 co-localizes with moesin at the apical membranes of kidney epithelial cells (**Figure 3D**). These data demonstrate that loss of TMIGD1 in mice interferes with the apical localization of the ERM proteins, moesin and ezrin. Moesin is highly enriched at the sites of cellular protrusions such as filopodia, microvilli and microspikes, and has been proposed to positively regulate cell migration in different cell types [8,11,31]. However, unlike moesin, TMIGD1 inhibits cell migration [2,10,27], suggesting a potential negative regulation of moesin function by TMIGD1. To address this question, we transfected HEK-293 cells expressing TMIGD1 or moesin with Life-ACT7 (GFP-fusion actin) and observed that HEK-293 cells expressing Life-ACT7 formed filopodia-like protrusions (**Figure 4A**), whereas HEK-293 cells expressing TMIGD1 with Life-ACT7 showed inhibition of filopodia-like protrusions (**Figure 4A**). Expression of TMIGD1 and moesin in HEK-293 is shown (**S. Figure 3**). In an additional strategy to probe the effect of TMIGD1 in filopodia protrusions, we expressed TMIGD1 in a colorectal carcinoma cell line, RKO cells and examined filopodia structures via phalloidin staining. We chose RKO cells because they form extensive filopodia-like protrusions. Expression of TMIGD1 in RKO cells significantly inhibited filopodia protrusions (**Figure 4B**). Expression of TMIGD1 in RKO cells is shown (**Figure 4B**). Furthermore, we expressed moesin-GFP in RKO cells and examined moesin localization in the filopodia structures. Moesin-GFP was mainly present in the filopodia structures in RKO cells (**Figure 4C**). However, the presence of moesin in filopodia protrusions in the context of TMIGD1 was less prominent (**Figure 4C**). Additionally, we analyzed the effect of expression of moesin in cell migration in RKO cells. The result showed that over-expression of moesin-GFP in RKO cells increased, whereas, expression of TMIGD1 inhibited cell migration (**Figure 4D**). Moreover, co-expression of TMIGD1 with moesin reversed the effect of moesin in cell migration (**Figure 4D**). Next, we knocked out moesin in RKO cells via CRISPR-Cas9 system and examined its effect in cell migration (**Figure 4E**). Loss of moesin in RKO cells significantly reduced cell migration in RKO cells and ectopic expression of TMIGD1 in these cells further decreased cell migration (**Figure 4F**). Taken together, these data demonstrate that moesin stimulates filopodia-like protrusions and cell migration and TMIGD1 inhibits its pro-migratory effects in HEK-293 and RKO cells.

**Figure 4.**
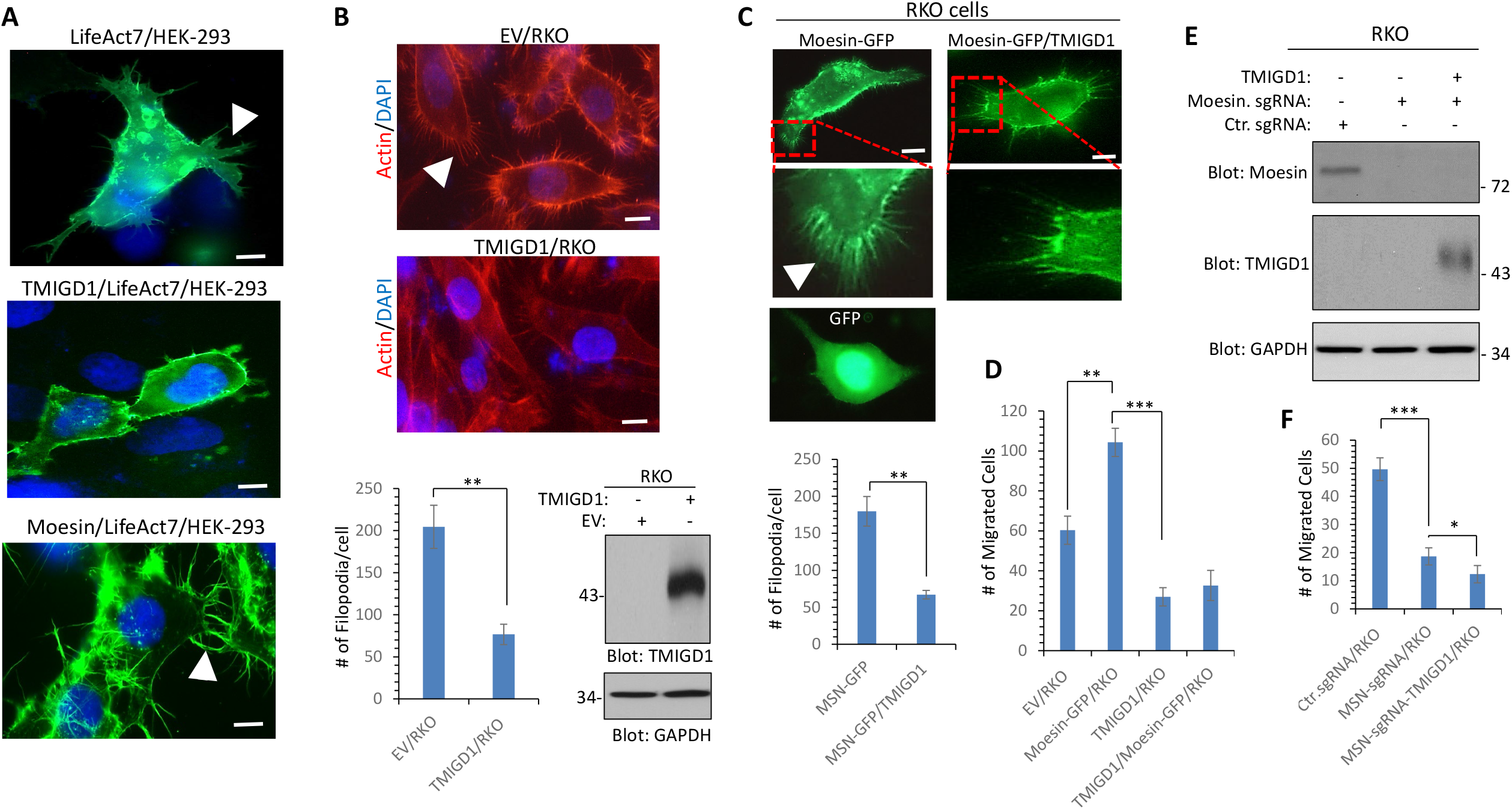
Moesin stimulates and TMIGD1 inhibits filopodia protrusions and cell migration. (**A**) HEK-293 cells and HEK-293 cells expressing TMIGD1 were transfected with Life-Act7. Filopodia protrusions observed under an epifluorescence microscope after 48hours transfection. Image magnification, 50 µM. White arrows point to filopodia-like protrusions. (**B**) RKO cells expressing empty vector or TMIGD1 were stained with phalloidin for actin (red) and DAPI (nucleus). Image magnification, 50µM. White arrow points to filopodia-like protrusions. The graph is a representative of filopodia formation in RKO cells expressing an empty vector (EV) or TMIGD1 (20 cells/group). **p<0.01. Cell lysates derived from RKO cells expressing empty vector or TMIGD1 were blotted for TMIGD1 and protein loading control, GAPDH. (**C**) RKO cells and RKO cells expressing TMIGD1 were transfected with Moesin-GFP. Moesin-GFP localization observed under an epifluorescence microscope after 48hours transfection. Image magnification, 50 µM. The graph is a representative of filopodia formation (based on the localization of moesin-GFP) in RKO cells expressing moesin-GFP (MSN-GFP) alone or co-expressing MSN-GFP with TMIGD1. **p<0.01. (**D**) The same cell groups were subjected to migration assay (quadruples/group) via Boyden chamber assay. **p<0.01. ***P <0.001. (**E**) Western blot analysis of CRISPR-Cas9 mediated knockout of moesin in RKO cells and RKO cells expressing TMIGD1. (**F**) Cells were subjected to cell migration assay (quadruples/group) via Boyden chamber assay system. *p<0.05, ***P <0.001.

### Expression of TMIGD1 in Renal Cancer Cells Promotes Stability of Microtubules

Among all the ERM family proteins (Ezrin, Radixin and Moesin), moesin is the only known member of ERM family proteins that has been shown to bind to microtubules and regulate their stability [37]. Considering the association of TMIGD1 with moesin, we decided to investigate whether TMIGD1 expression in renal cell carcinoma, 786-0 cell line could modulate microtubule stability. First, we examined whether TMIGD1 can affect Lysine 40 (K40) acetylation of tubulin in 786-0 cells, because K40 acetylation is critically important for microtubular dynamics and stability [43]. Immunofluorescence staining with an antibody against K40-acetylated tubulin in 786-0 cells expressing empty vector (EV) or TMIGD1 revealed that the level of K40-acetylated tubulin in 786-0 cells expressing TMIGD1 was significantly higher than observed for the cells expressing EV (**Figure 5A**). Western blotting analysis further validated the data obtained from immunofluorescence staining analysis (**Figure 5B**). Next, we asked whether expression of TMIGD1 in 786-0 cells could prevent microtubule depolymerization induced by nocodazole, a small molecule that binds to tubulin and thereby inhibits microtubule depolymerization [20,24]. Our result showed that expression of TMIGD1 in 786-0 cells prevented nocodazole-induced microtubule depolymerization (**Figure 5C**). The effect of TMIGD1 was quantified at various times after nocodazole treatment by counting the number of cells displaying intact microtubules. The most striking effect of TMIGD1 on the microtubule stability was observed at 20-30 min after nocodazole treatment (**Figure 5C**). Representative images of the effect of nocodazole on RKO cells at 20 and 30 minutes are shown (**S. Figure 4**). We further investigated the role of TMIGD1 in the microtubule mitotic spindle organization. Consistent with the observed effect of TMIGD1 stabilizing microtubules, 786-0 cells expressing TMIGD1 undergoing mitosis developed well-formed microtubule mitotic spindles, whereas the formation of spindles in EV/786-0 cells were relatively short and less organized (**Figure 5D**). Interestingly, there were also markedly high levels of actin on the periphery of 786-0 cells expressing TMIGD1 (**Figure 5D**).

**Figure 5:**
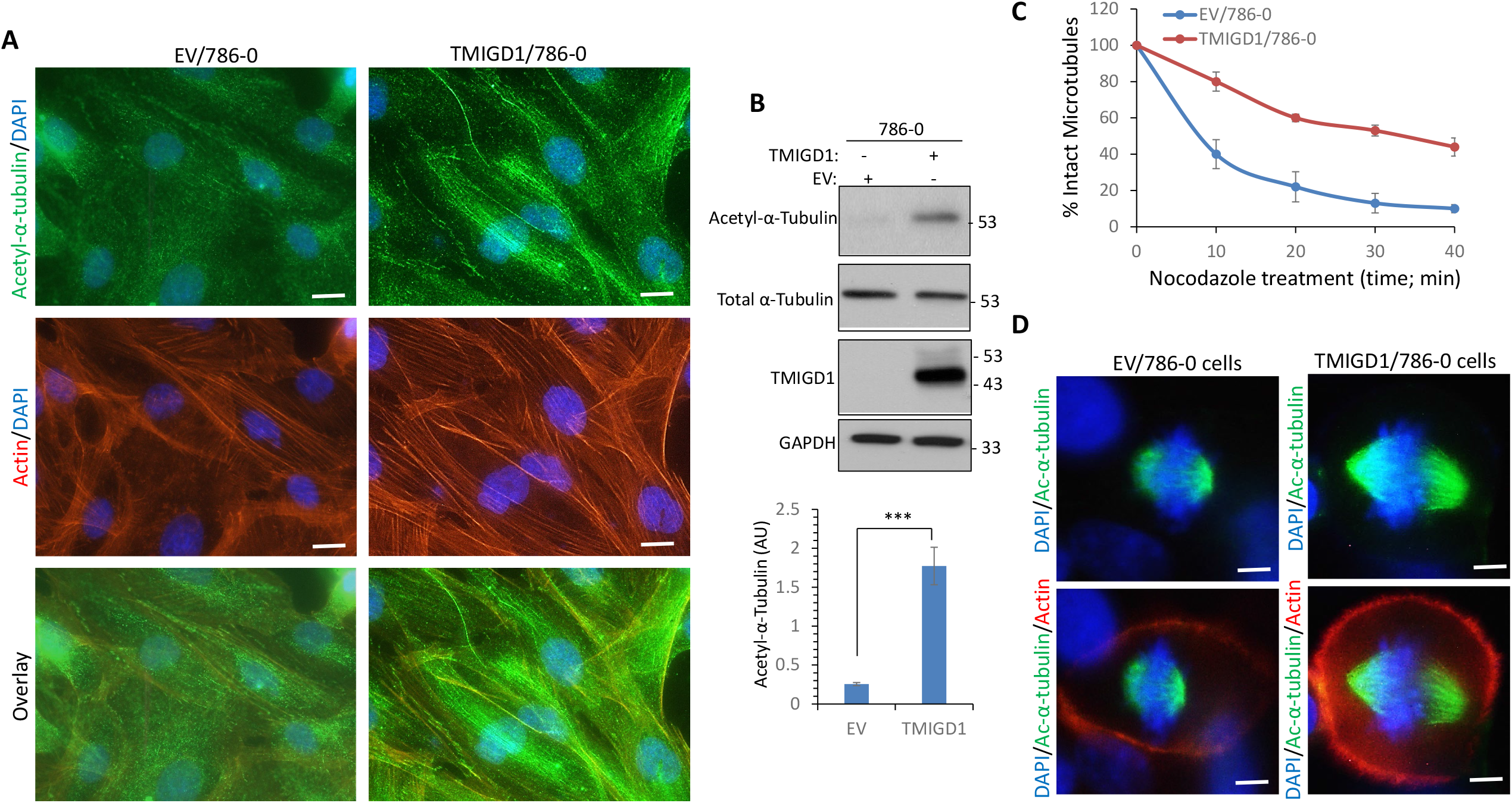
TMIGD1 stimulates α-tubulin acetylation and regulates mitotic spindle assembly. (**A**) EV/786-0 and TMIGD1/786-0 cells were stained with an antibody to K40 acetyl-α-tubulin (green) and phalloidin/actin (red). (**B**) Cell lysates from the same cell groups were blotted for acetylated-α-tubulin, total α-tubulin and TMIGD1. The graph represents three independent experiments.***P <0.001. (**C**) EV/786-0 and TMIGD1/786-0 cells (triplicate per group) were treated with nocodazole (0.5µg/ml) for various times, followed by staining with an antibody against acetylated-α-tubulin. The numbers of cells displaying intact microtubules were manually counted (n = 50 cells per each group of triplicates). (**D**) EV/786-0 and TMIGD1/786-0 cells were stained with acetylated-α-tubulin (green) and phalloidin/actin (red) and the pictures of mitotic cells (n = 20 cells per group) were taken under an epifluorescence microscope.

Following this finding, we asked whether inactivation of moesin via a CRISPR-Cas9 system in 786-0 cells interferes with the TMIGD1-dependent acetylation of tubulin, which is critical for microtubule mitotic spindle organization (**Figure 6A**). Knockout of moesin in 786-0 cells inhibited basal acetylation (K40) of tubulin and, more importantly, markedly blocked TMIGD1-dependent acetylation of tubulin (**Figure 6A**). Similar effects were observed via immunofluorescence staining with anti-K40-acetyl-tubulin antibody (**Figure 6B**). Considering the roles of both TMIGD1 and moesin in actin fibril organization, we examined whether the loss of moesin also affects actin fibril formation. The result showed that loss of moesin inhibited TMIGD-1 mediated F-actin orientation (**Figure 6C**). Taken together, the data demonstrate that TMIGD1 dependent K40 acetylation of tubulin and F-actin orientation are mostly mediated by moesin.

**Figure 6.**
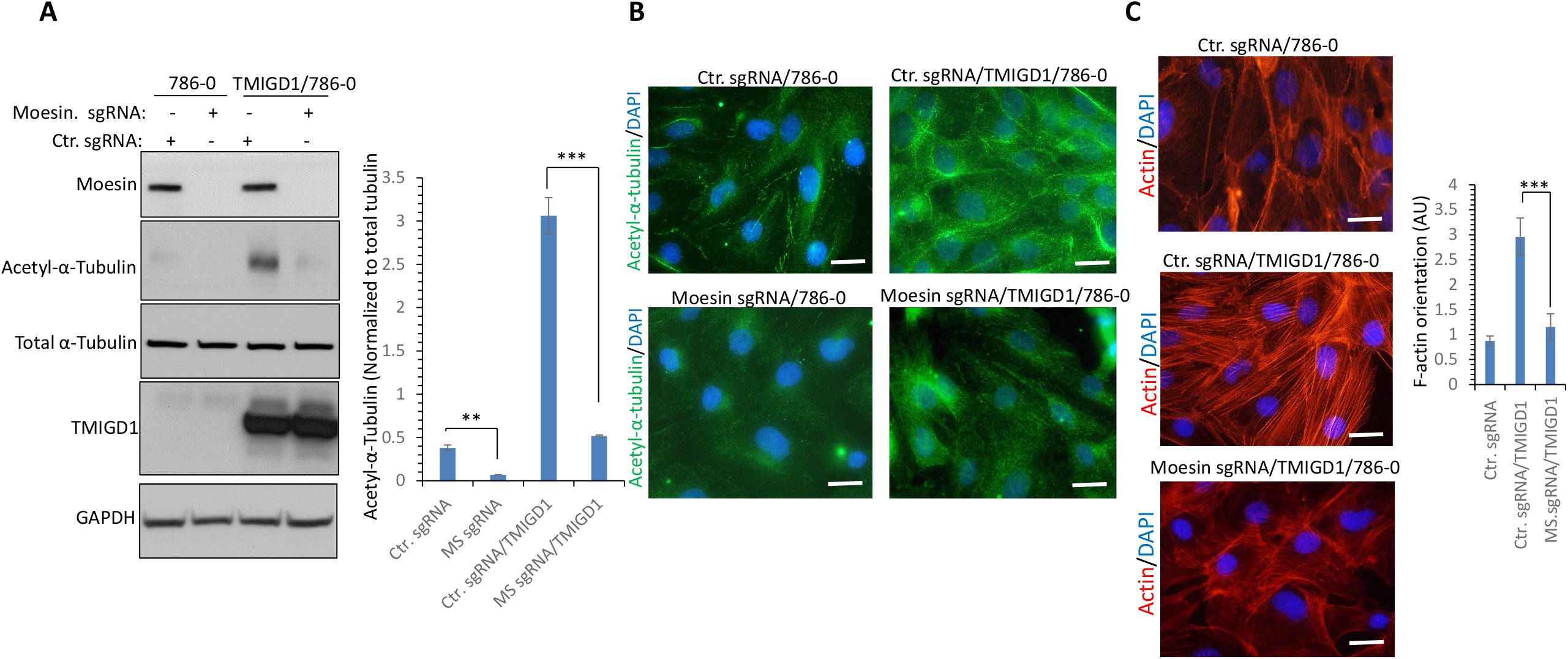
Knockout of Moesin in 786-0 cells inhibits TMIGD1-dependent α-tubulin acetylation and F-actin orientation. (**A**) Whole cell lysates from 786-0 cells expressing control sgRNA or moesin-sgRNA (MS-sgRNA) and 786-0 cells co-expressing control sgRNA with TMIGD1 or MS-sgRNA with TMIGD1 were subjected to Western blot via blotting for moesin, acetyl-tubulin, total tubulin, TMIGD1 or GAPDH. ***P <0.001, **p<0.01. (**B**) The same cell lines as panel A were subjected to immunofluorescence staining using anti-acetyl-tubulin antibody. Image magnification, 50µM. 786-0 cells expressing Ctr. sgRNA, Ctr.sgRNA/TMIGD1 or moesin-sgRNA/TMIGD1 were stained with phalloidin/DAPI and actin fibril orientation/anisotropy was quantified. (**C**) EV/786-0 and TMIGD1/786-0 cells were treated with nocodazole (0.5µg/ml) for various times followed by staining with an antibody against acetylated-α-tubulin. The numbers of cells displaying intact microtubules were manually counted (n = 50 cells per each group of triplicates). EV/786-0 and TMIGD1/786-0 cells were stained with an antibody against acetylated-α-tubulin (green) and phalloidin/actin (red) and pictures of mitotic cells (n = 20 cells per group) were taken under an epifluorescence microscope.

### High expression of TMIGD1, Moesin and Ezrin correlates with better renal cell carcinoma survival

Having established a functional link between TMIGD1 and moesin, we asked whether expression profiles of the TMIGD1 and ERM family proteins, moesin and ezrin, correlate with survival of renal cell carcinoma (RCC) patients. Our previous analysis of data sets from The Cancer Genome Atlas and our own immunohistochemical analysis revealed that low expression of TMIGD1 correlates with poor survival in renal and colon cancers[10,27]. Although, moesin and ezrin had been proposed to function as oncogenes in various tumor cell lines [8], their potential roles in RCC is not known. To address this question, we carried out a Kaplan-Meier survival analysis of the data set for clear cell renal cell carcinoma, the most common form of RCC, which is publicly available from The Cancer Genome Atlas via the Kaplan-Meier Plotter[29]. The result revealed that patients with RCC and high TMIGD1-, moesin- or ezrin-expressing tumors had significantly better median overall survival compared with those with low TMIGD1-expressing tumors (**Figure7A-C**). The median survival for patients with tumors expressing high TMIGD1 (n=323) was 118.47 months versus 66 months for tumors with low TMIGD1 expression (n=207) (**Figure 7A**). Patients with high moesin- and ezrin-expressing tumors also showed significantly better survival (mean survival time for patients with tumors expressing high moesin was 52.8 months versus 27.3 months for those with low expression; the mean survival time for patients with tumors expressing high ezrin was 66.2 months versus 27.4 months for those with low expression) (**Figure 7B, C**).

**Figure 7.**
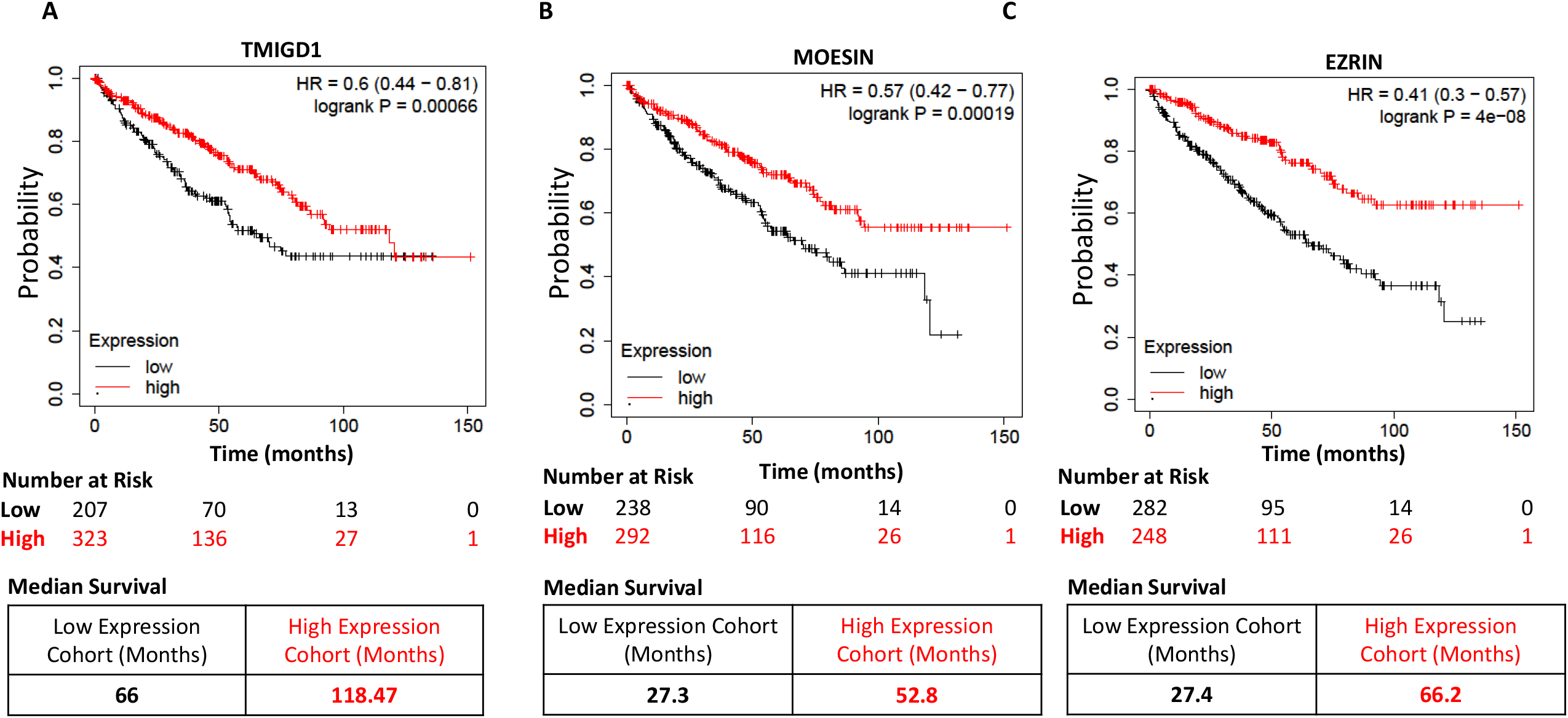
High TMIGD1, Moesin and Ezrin expression in renal colon cancer correlate with better survival. Kaplan Meier survival analysis of kidney renal clear cell carcinoma for high and low expression of TMIGD1, moesin and ezrin. Total number of cases 530. Lower panel tables show median survival (**A-C**). The data derived from the Cancer Genome Atlas (TCGA) data set via Kaplan Meier plotter [29].

Having observed a correlation between better survival rate and high moesin and ezrin expression levels in human RCC, we asked whether moesin and ezrin mRNA levels are altered in RCC using the same TCGA data set via Fire Brows (http://firebrowse.org/). Our analysis demonstrated that TMIGD1 mRNA is downregulated in all the major kidney cancer types including, kidney clear cell renal cell carcinoma (KIRC), kidney papillary renal cell carcinoma (KIRP) and chromophobe renal cell carcinoma (KICH) (**S. Figure 4A**). The median expression of TMIGD1 mRNA in KIRC was 2.32 versus 3.62 normal tissues, in KIRP was 1.16 versus 2.44 and in KICH was -1.3 versus 1.191 normal tissue (**S. Figure 5A**). The median expression of TMIGD1 mRNA in pan kidney cancers (KIPAN) was 1.91 versus 3.42 normal (**S. Figure 5A**). Interestingly, the expression levels of neither moesin nor ezrin were altered in a statistically significant manner in any of the kidney cancer types (**S Figure 5B, C**). The median expression of moesin mRNA in KIRC was 14.2 versus 13.7 normal tissues, in KIRP was 13.4 versus 13.3 and in KICH was 12.1 versus 13.4 normal tissue (**S. Figure 5B**). The median expression of moesin mRNA in KIPAN was 13.6 versus 13.9 normal (**S. Figure 5B**). Similarly, the median expression of ezrin mRNA in KIRC was 13.7 versus 13.4 normal tissues, in KIRP was 14.5 versus 13.5 and in KICH was 12.1 versus 13.4 normal tissue (**S. Figure 5C**). The median expression of ezrin mRNA in KIPAN was 12.4 versus 13.4 normal (**S. Figure 5C**).

Considering that we did not observe any major changes in the mRNA levels of moesin and ezrin in the human kidney cancers, we decided to analyze the TCGA data set for the presence of potential mutations on moesin and ezrin. We found no mutations on moesin or ezrin; only two or three cases of missense mutations were found (data not shown). Similarly, no significant mutations were found on MIGD1 in RCC. There were six cases of missense and three truncating mutations. The somatic mutation frequency of TMIGD1 was 0.3% (Data not shown). Our analysis of the TCGA data demonstrating the low frequency of mutations on moesin and ezrin combined with no major alterations in their mRNA levels suggests that deregulations of cellular localization or posttranslational modifications of moesin and ezrin could account for their altered roles in tumorigenesis. Expression of TMIGD1 both at the protein levels as previously reported [10,27] and at the mRNA levels (**S. Figure 5A**) is downregulated in RCC indicating that downregulation of TMIGD1 is a primary mechanism of its alteration in human RCC which may also account for deregulation of downstream signaling mechanisms such as signaling by the ERM family proteins moesin and ezrin in human cancer cells.

## Discussion

The cell adhesion molecule TMIGD1 is newly identified tumor suppressor that regulates critical cellular processes such as cell-cell adhesion, cell migration, cell proliferation and cell cycle[2,10,27]. However, the molecular mechanisms governing its signaling mechanism remains largely unknown. In this study, we identified moesin as a major TMIGD1 binding protein and a key regulator of α-tubulin acetylation. TMIGD1 recognizes and associates with the N-terminal FERM domain of moesin via the conserved RRKK motif located on the cytoplasmic domain and regulates moesin cellular localization. Loss of TMIGD1 in mice impairs apical membrane localization of moesin and ezrin in the epithelium of renal and intestinal tissues, respectively; cell culture studies reveal that moesin stimulates the formation of filopodia protrusions and promotes cell migration. However, co-expression of moesin with TMIGD1 hinders the pro-migratory effects of moesin. Furthermore, moesin has been proposed to regulate various cellular processes such as cell migration that are considered pro-invasion and pro-metastatic[8], whereas our study suggests that the role of moesin in these cellular processes is more nuanced and rather context-dependent. For example, the expression status of TMIGD1 in human cancers, such as colon and renal cancers, could determine whether moesin and ezrin function as pro- or anti-tumorigenic factors. Furthermore, our study suggests that loss of TMIGD1 in human cancers, which was previously reported in renal and colon cancers [5,10,27], could result in the activation of pro-tumorigenic signaling cascades, including the activation of ERM proteins that contribute to tumor progression.

Another important and interesting aspect of our study is the finding that TMIGD1 appears to diminish the pro-tumorigenic properties (e.g., cell migration and acetylation of tubulin) of moesin in cells perhaps through the process of binding and sequestering moesin to the apical domain of epithelial cells. Recruitment of moesin to TMIGD1 mediates microtubular regulation, specifically aiding the K40 acetylation of α-tubulin. The acetyltransferase aTAT1 is responsible for the K40 acetylation of α-tubulin [1,19] and tubulin acetylation has been shown to stimulate cell migration in multiple cell types including, fibroblasts [17], neuronal [9] and astrocytes [3], suggesting that TMIGD1/moesin pathway could regulate cell migration via modulating tubulin acetylation. Previous studies have shown that moesin connects major cytoskeletal structures to the plasma membrane and regulates cell polarity, cell adhesion and microtubule dynamics [7,35,36]. Since the dynamics of microtubules are altered in cancer cells and are linked to chromosomal instability and development of drug resistance [6],[18], they are considered an attractive target for chemotherapy against various cancer types[16]. Moesin binds directly to microtubules and regulates spindle organization in metaphase, cell morphogenesis during mitosis and cell polarity [25,36,38], suggesting that, by interacting with TMIGD1, Moesin links TMIGD1 to microtubules and thus modulates cell polarity.

## Conclusions

Identification of TMIGD1 as an upstream receptor capable of regulating the activity of ERM family proteins offers new insights into the mechanisms of TMIGD1 tumor suppressor signaling and tumorigenesis. We show for the first time that TMIGD1 binds to ERM family proteins, regulates the stability of microtubules and modulates cell migration in renal cancer cells, implicating TMIGD1 as an attractive potential cancer therapeutic target. Further investigation into mechanisms by which TMIGD1 regulates moesin cellular localization and activity should provide an increasingly mechanistic explanation of the interplay between TMIGD1 and ERM family proteins. Furthermore, considering our observation that higher expression profiles of TMIGD1, moesin and ezrin correlate with better survival, deeper investigation into the molecular basis of downregulation of TMIGD1, moesin and ezrin will shed new insight into the altered roles of cell adhesion signaling mechanisms in tumor progression.

## Declarations

### Ethics approval and consent to participate

The care and use of laboratory animals were approved by Boston University.

### Consent for publication

All authors approve the publication of the manuscript.

### Availability of data and materials

Cell lines, plasmids and other reagents described in this manuscript are available upon a reasonable request.

## Authors’ Contribution

NR, CEC and RX-YO were involved in writing and editing of the manuscript. NR, KBC, KCSDC, RA, AM and NE all were involved in the design, performing the experiments.

## Acknowledgements

Not applicable.

