## supplemental data for "The Cell Adhesion Molecule TMIGD1 Binds to Moesin and Regulates Tubulin Acetylation and Cell Migration"

**S. Figure 1**: Amino acid sequence homology of human moesin and ezrin. Alignment of human moesin (MSN) and ezrin (EZN) is shown. The FERM domain is shown in a box. Hydrophobic and negatively amino acids are highlighted in light blue and light brown colors, respectively.

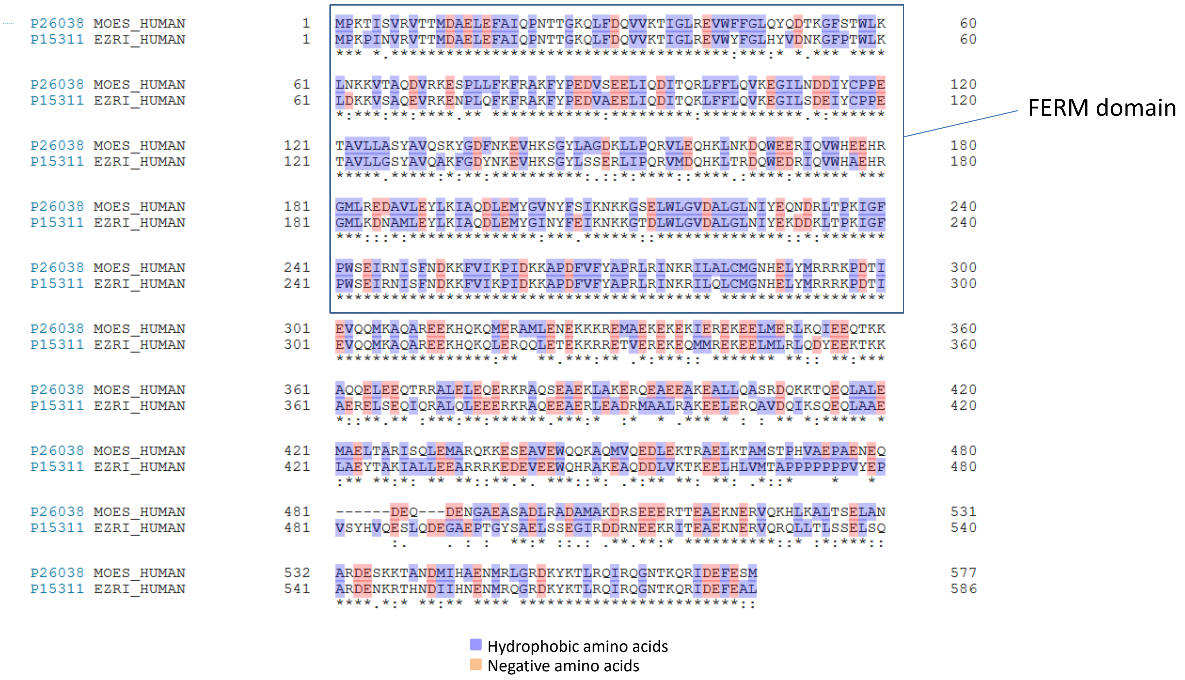


**S. Figure 2. qPCR analysis of TMIGD1 KO mouse:** qPCR analysis of wild-type and TMIGD1 is shown.

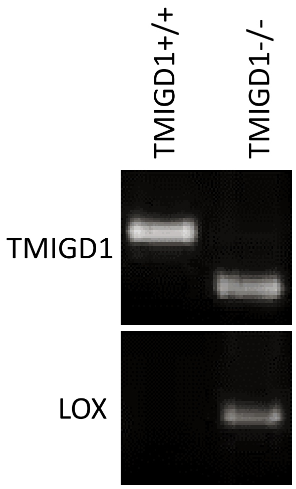

**S. Figure 3. Ectopic expression of Moesin and TMIGD1 in HEK-293 cell**. Whole cell lysates derived from HEK-293 cells expressing TMIGD1 or moesin subjected to western blot analysis.

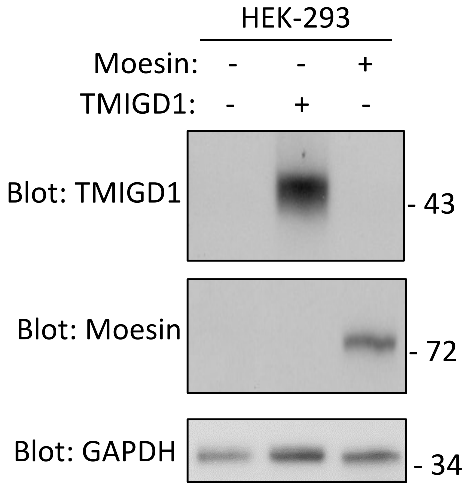

**S. Figure 4. Effect of Nocodazole in the microtubules of RKO cells expressing TMIGD1.** EV/786-0 and TMIGD1/786-0 cells (triplicate per group) were treated with nocodazole (0.5µg/ml) for 20 or 30 minutes, followed by staining with an antibody against acetylated-α-tubulin.

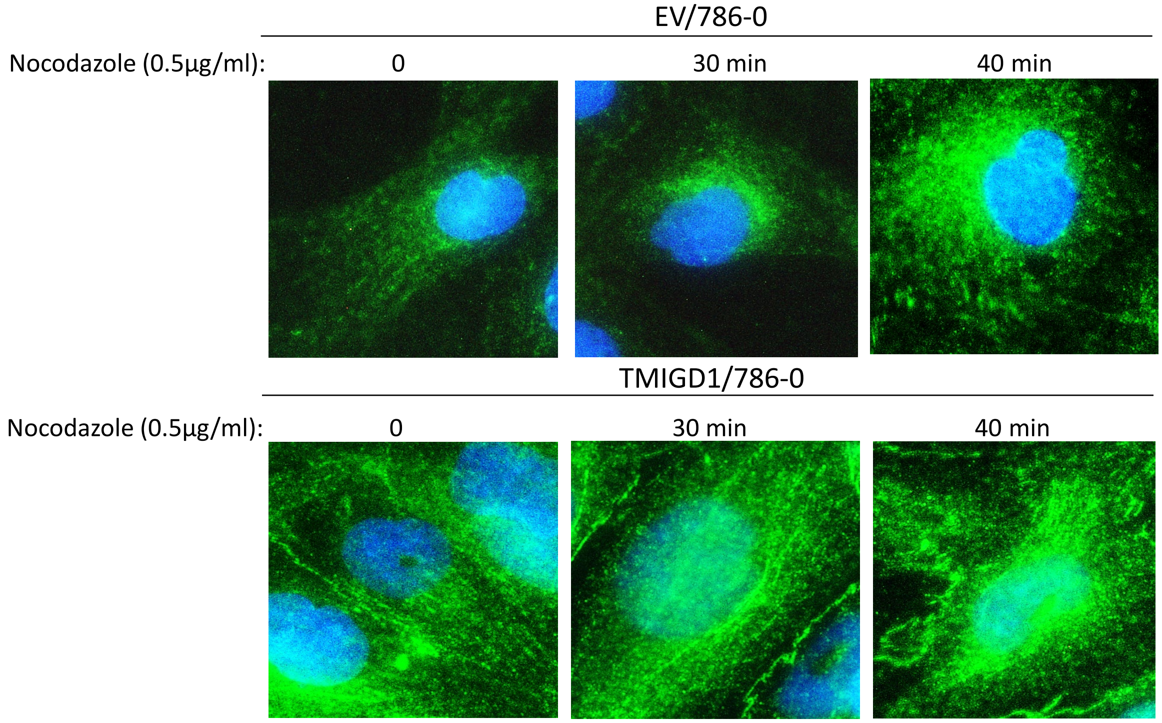

**S. Figure 5**: **TMIGD1, Moesin and Ezrin mRNA levels in human normal kidney verses kidney cancer types**. (**A-C**) TMIGD1, Moesin and Ezrin mRNA levels obtained from the TCGA data set via Fire Brows (http://firebrowse.org/). Kidney cancer types; kidney clear cell renal cell carcinoma (KIRC), kidney papillary renal cell carcinoma (KIRP), chromophobe renal cell carcinoma (KICH) and kidney pan cancer analysis (KIPAN). The horizontal small black bars within each box correspond to median expression of given proteins.

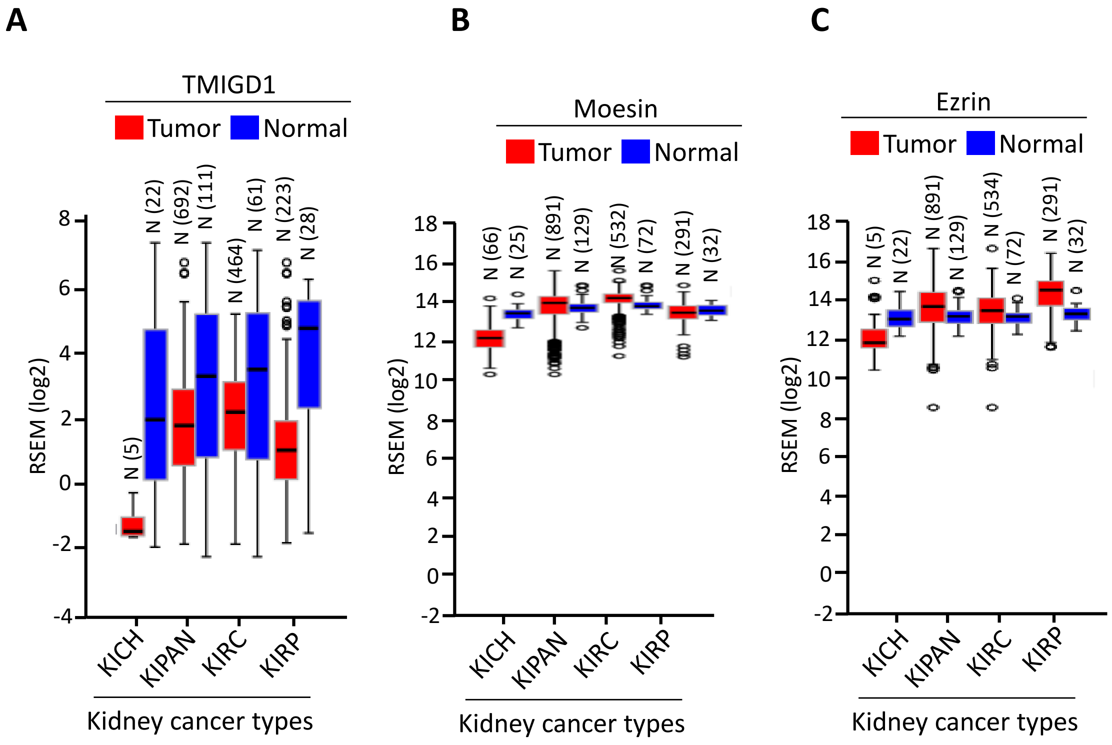
